# Targeting MCL1-driven anti-apoptotic pathways to overcome hypomethylating agent resistance in *RAS*-mutated chronic myelomonocytic leukemia

**DOI:** 10.1101/2023.04.07.535928

**Authors:** Guillermo Montalban-Bravo, Feiyang Ma, Natthakan Thongon, Hui Yang, Irene Ganan- Gomez, Juanjo Jose Rodriguez-Sevilla, Vera Adema, Bethany Wildeman, Pamela Lockyer, Yi June Kim, Tomoyuki Tanaka, Faezeh Darbaniyan, Shivam Pancholy, Geoffrey Zhang, Gheath Al-Atrash, Karen Dwyer, Koichi Takahashi, Guillermo Garcia-Manero, Hagop Kantarjian, Simona Colla

## Abstract

*RAS* pathway mutations, which are present in 30% of patients with chronic myelomonocytic leukemia (CMML) at diagnosis, confer a high risk of resistance to and progression after hypomethylating agent (HMA) therapy, the current standard of care for the disease. Using single-cell, multi-omics technologies, we sought to dissect the biological mechanisms underlying the initiation and progression of *RAS* pathway–mutated CMML. We found that *RAS* pathway mutations induced the transcriptional reprogramming of hematopoietic stem and progenitor cells (HSPCs), which underwent proliferation and monocytic differentiation in response to cell-intrinsic and -extrinsic inflammatory signaling that also impaired immune cells’ functions. HSPCs expanded at disease progression and relied on the NF-_K_B pathway effector MCL1 to maintain their survival, which explains why patients with *RAS* pathway– mutated CMML do not benefit from BCL2 inhibitors such as venetoclax. Our study has implications for developing therapies to improve the survival of patients with *RAS* pathway– mutated CMML.

## INTRODUCTION

Chronic myelomonocytic leukemia (CMML), a clonal disorder of mutant hematopoietic stem cells (HSCs)^1^, is characterized by myelodysplastic and myeloproliferative bone marrow (BM) features^2,3^ and a high risk of progression to acute myeloid leukemia (AML)^4-6^. Although hypomethylating agent (HMA) therapy, the current standard of care for most patients with CMML^7^, can overcome CMML cells’ aberrant proliferation and achieve improved outcomes in some patients, most patients have only transient responses to these agents, owing to the agents’ inability to effectively deplete HSCs and decrease tumor burden. CMML patients whose disease undergoes transformation to AML upon HMA therapy failure have dismal clinical outcomes^8^.

Despite advances in the genetic characterization of CMML, the development of alternative frontline treatments or more effective second-line therapies to improve the outcomes of CMML patients with high-risk biological features has been delayed because of an incomplete understanding of the ways in which different hematopoietic populations that persist throughout HMA therapy contribute to disease maintenance and progression.

Mutations in *RAS* pathway signaling genes (*BRAF, CBL, KRAS, NF1, NRAS,* and *PTPN11*) confer adverse biological features that increase the risk of disease progression and poor overall survival, particularly when they are present with loss-of-function mutations in the *ASXL* transcriptional regulator 1 gene, *ASXL1*^9^.

Here, we used single-cell technology–based approaches to elucidate the biological and molecular landscape of *RAS* pathway–mutated CMML to guide the selection of novel therapeutic interventions and achieve lasting responses in CMML patients in whom HMA therapy has failed.

## RESULTS

### Mutations in *RAS* pathway signaling genes predict a high risk of CMML blast progression after HMA therapy failure

We first evaluated whether specific mutations predict a high risk of CMML blast progression (BP) in a cohort of 108 CMML patients who received HMA therapy (Supplementary Table S1). After a median follow-up of 19 months (95% confidence interval [CI], 15.8–23.9 months), 57 patients had experienced HMA therapy failure. Among 36 patients who had increased blast counts at the time of therapy failure, 13 had progression to AML. Mutations in *RAS* pathway genes were associated with BP (odds ratio, 3.35; 95% CI, 1.46–7.70; *P*=0.004) (Fig. 1A) and shorter time to BP (hazard ratio, 2.21; 95% CI, 1.13–4.33; *P*=0.021) (Fig. 1B). Logistic regression analysis similarly showed that *RAS* pathway mutations were associated with the risk of BP (*P*=0.01158) (Supplementary Table S2). To assess whether BP was associated with mutations that were not detected at diagnosis or with the clonal expansion of pre-existing mutations, we sequenced BM cells isolated from samples collected at the time of HMA therapy failure from 22 of the 36 patients with BP and compared the cells’ genomic landscape with that of BM cells isolated at diagnosis (Fig. 1C). Among the 22 patients with BP, 14 (64%) had *RAS* pathway mutations at diagnosis, 20 (91%) had *RAS* pathway mutations at BP, and 9 (41%) had acquired newly detectable *RAS* pathway mutations at BP (4 of these patients had no detectable *RAS* pathway mutations at diagnosis and 5 acquired other *RAS* pathway mutations). Of the 14 patients with *RAS* pathway–mutated CMML at diagnosis, 10 had BP without *RAS* pathway mutation–induced clonal evolution (Fig. 1D).

**Figure 1.**
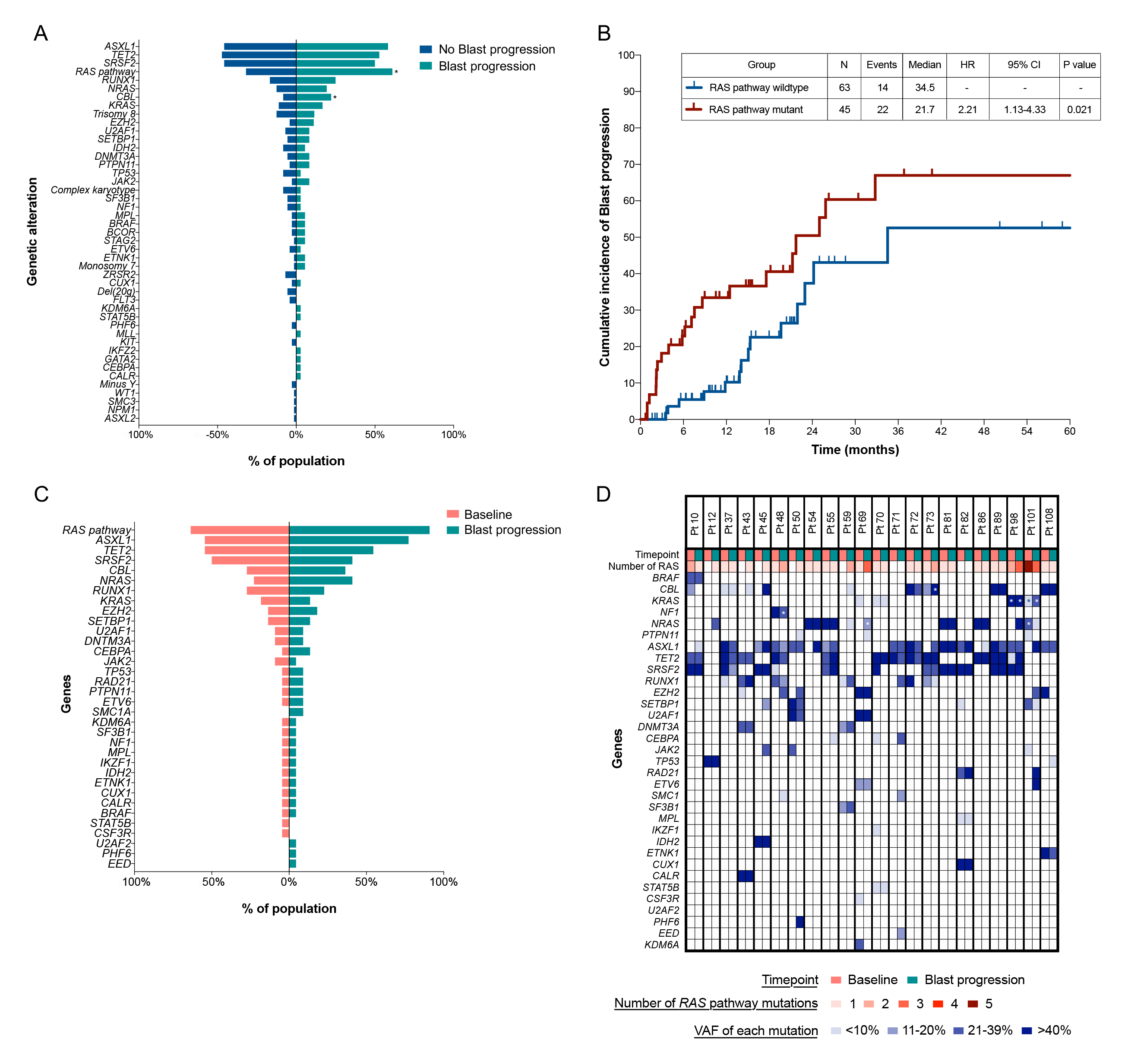
Mutations in *RAS* pathway signaling genes predict a high risk of CMML BP after HMA therapy failure. (A) Bar chart showing the frequencies of detectable mutations and cytogenetic abnormalities among 108 CMML patients who received HMA therapy and whose disease progressed (green) or did not progress (blue). Asterisks indicate significantly different frequency changes (\**P*<0.05). (B) Kaplan-Meier survival curves showing the cumulative incidence of BP after HMA therapy in previously untreated CMML patients with or without *RAS* pathway mutations. N, number; HR, hazard ratio; CI, confidence interval. (C) Bar chart showing the overall frequencies of detectable mutations among 22 CMML patients whose disease progressed and in whom targeted sequencing was performed at the time of BP. Mutations at diagnosis and BP are indicated by pink and green, respectively. Paired samples were analyzed. (D) Detected mutations and their variant allele frequencies (VAFs) in matched samples obtained at diagnosis and at the time of BP in the 22 CMML patients shown in Fig. 1C. Columns represents the mutations and VAFs of individual CMML patients’ sequential samples at diagnosis and BP. Patient identifiers are shown at the top of each column. Asterisks indicate the presence of multiple mutations for a particular gene. The numbers of *RAS* mutations are shown in red gradient; the VAFs of each mutation are shown in blue gradient.

These results were validated using single-cell DNA sequencing coupled with cell-surface immunophenotyping analysis of mononuclear cells (MNCs) isolated from sequential BM samples obtained at the time of diagnosis or BP from 2 representative *RAS* pathway– mutated CMML patients whose disease never responded to therapy (Supplementary Fig. S1A, S1B) or underwent clonal evolution after an initial response (Supplementary Fig. S1C, S1D). Together, these data demonstrate that patients with *RAS* pathway–mutated CMML have a high risk of BP at the time of HMA therapy failure. This observation has important clinical implications in light of our recent study showing that *RAS* pathway mutations also drive resistance to and/or BP following venetoclax-based second-line therapy^10^. These data underscore the urgent need to dissect the biological mechanisms of *RAS* pathway mutation– induced therapy resistance, as such an understanding could lead to the development of new therapeutic approaches to prevent or overcome disease progression.

### *RAS* pathway–mutations activate cell-intrinsic and -extrinsic inflammatory networks

To dissect the molecular mechanisms underlying the progression of *RAS* pathway– mutated CMML, we first evaluated the molecular determinants of the disease initiation. We performed single-cell RNA sequencing (scRNA-seq) analysis of lineage-negative (Lin^−^) CD34^+^ hematopoietic stem and progenitor cells (HSPCs) isolated from 5 untreated CMML patients and 2 age-matched healthy donors (HDs) (Supplementary Table S3). This analysis identified 8 cellular clusters driven by the cells’ differentiation profile (Fig. 2A) that we defined based on the differential expression of validated lineage-specific transcriptional factors (TFs) and cellular markers^11,12^ (Supplementary Fig. S2A; Supplementary Table S4). Compared with those from HDs, Lin^−^CD34^+^ HSPCs from CMML patients had a predominant myeloid differentiation route with higher frequencies of early myeloid hematopoietic progenitor cells (eMyHPCs; clusters 1 and 3, characterized by the high expression of *CD34*, *BTF3*, and *CEBPA* but low expression of *CD38*) and more differentiated MyHPCs (dMyHPCs; clusters 0, 4 and 6, marked by the expression of *CEBPD* and/or *CEBPA*, as well as that of *MPO*) at the expense of more primitive HSCs (cluster 2, marked by the high expression of *MLLT3*, *MEG3*, and *CLEC9A*) and erythroid/megakaryocyte (Ery/Mk) HPCs (clusters 5 and 7, marked by the expression of *KLF1*, *GATA1*, and *GATA2*) (Fig. 2B). Differential expression analysis revealed that genes upregulated in CMML HSCs compared with HD HSCs were mainly involved in oxidative phosphorylation, interferon (IFN) response, and apoptosis (Fig. 2C; Supplementary Fig. S2B). Similar results were observed in eMyHPCs and dMyHPCs (Fig. 2C).

**Figure 2.**
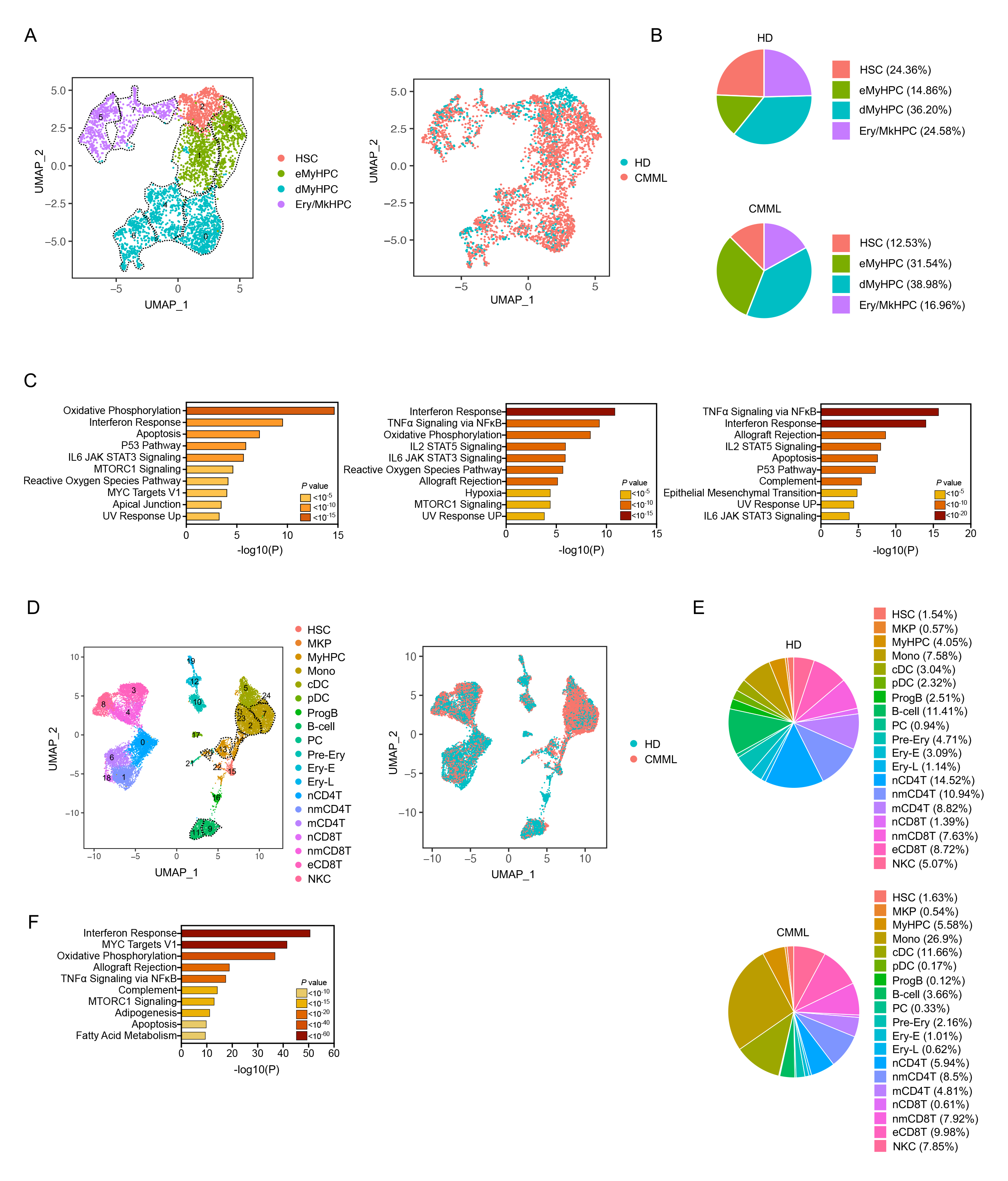
*RAS* pathway–mutated CMML cells activate cell-intrinsic and -extrinsic inflammatory networks. (A) UMAP of scRNA-seq data for pooled single Lin^−^CD34^+^ cells isolated from BM samples of 2 HDs (n=895) and 5 CMML patients (n=3,161). Each dot represents one cell. Different colors represent the cluster cell type identity (left) or sample origin (right). HSC, hematopoietic stem cells; eMyHPC, early myeloid progenitor cells; dMyHPC, differentiated myeloid progenitors; Ery/MkHPC, erythroid/megakaryocyte hematopoietic progenitor cells. Dashed lines indicate single clusters in each cell type population. (B) Distribution of HD (top) and CMML (bottom) Lin^−^CD34^+^ cell types among the clusters shown in Fig. 2A. (C) Pathway enrichment analysis of the genes that were significantly upregulated in HSCs (left), eMyHPCs (middle), and dMyHPCs (right) from CMML samples compared with those from HD samples (adjusted *P* ≤ 0.05). The top 10 Hallmark gene sets are shown. (D) UMAP of scRNA-seq data for pooled single MNCs isolated from BM samples of 3 HDs (n=9,896) and 5 CMML patients (n=9,319). Each dot represents one cell. Different colors represent the cluster cell type identity (left) or sample origin (right). HSC, hematopoietic stem cells; MKP, megakaryocyte precursors; MyHPC, myeloid hematopoietic progenitor cells; Mono, monocytes; cDC, classical dendritic cells; pDC, plasmacytoid dendritic cells; Prog B, progenitor B-cells; PC, plasma cells; Pre-Ery, pre-erythrocytes; Ery-E, early erythroid precursors; Ery-L, late erythroid precursors; nCD4T, naïve CD4^+^ T cells; nmCD4T, naïve and memory CD4^+^ T cells; mCD4T, memory CD4^+^ T cells; nCD8T, naïve CD8^+^ T cells; nmCD8T, naïve and memory CD8^+^ T cells; eCD8T, effector CD8^+^ T cells; NKC, natural killer cells. Dashed lines indicate single clusters in each cell type population. (E) Distribution of HD (top) and CMML (bottom) cell types among the clusters shown in Fig. 2D. (F) Pathway enrichment analysis of the genes that were significantly upregulated in the CMML monocyte clusters compared with those in the HD monocyte clusters shown in Fig. 2D (adjusted *P* ≤ 0.05). The top 10 Hallmark gene sets are shown.

To evaluate the contribution of downstream myelo/monocytic (My/Mo) populations to disease maintenance, we performed scRNA-seq analysis of MNCs isolated from 3 HDs’ and 5 untreated CMML patients’ BM samples that were previously analyzed at the HSPC level. This analysis identified 25 cellular clusters inclusive of all major BM cell types that we defined based on the expression of lineage-specific TFs and cellular markers and using the single cell Transcriptome to Protein prediction with deep neural network (cTP-net) pipeline^13,14^ (Fig. 2D; Supplementary Fig. S2C; Supplementary Table S5). Consistent with the predominant myeloid differentiation bias of CMML HSPCs, differential analysis of BM cell lineage composition revealed that the monocyte population (clusters 2, 7, 23, and 24) was significantly higher in CMML BM samples than in HD BM samples (Fig. 2E). CMML monocytes underwent significant transcriptional reprogramming and mainly overexpressed genes involved in inflammatory signaling responses mediated by IFN and the NF-_K_B pathway (Fig. 2F; Supplementary Fig. S2D). Cytokines encoded by *S100A* genes, including *S100A8*, *S100A9*, and *S100A12,* and these cytokines’ receptor, *TLR4*, whose interactions further activate NF-_K_B pathway–induced inflammatory cascade, were also significantly upregulated in CMML monocytes (Supplementary Fig. S2E). Notably, IFNγ and TNFα, whose expression was not upregulated in CMML monocytes, were among the predicted upstream regulators of this transcriptional signature (Supplementary Fig. S2F), which suggests that other extrinsic cytokines contribute to activating the inflammasome response. Together, these data are consistent with previous findings in other cancer types that NF-_K_B signaling–induced inflammasome activation is essential for *RAS* pathway mutation–induced tumorigenesis^15-18^.

Inflammatory networks modulate the immune microenvironment and contribute to immune escape^19^. To assess whether *RAS* pathway–mutated CMML monocytes directly suppress the immune response, we dissected the inter-cellular crosstalk and communication networks between CMML cells and all other BM cells. We inferred cell-to-cell communication from the combined expression of multi-subunit ligand–receptor complexes using CellPhoneDB, a repository of ligands and receptors and their interactions ^20^. After generating a homeostatic interactome of BM MNCs from HDs, we analyzed the cellular communication networks upregulated in CMML BM samples (Supplementary Fig. S2G; Supplementary Table S6). Compared with HD MNCs, CMML MNCs had significantly more ligand–receptor interactions involving monocytes, classical dendritic cells (cDCs), plasmacytoid DCs (pDCs), MyHPCs, effector CD8^+^ T (eCD8T) cells, and natural killer (NK) cells. Specifically, chemokine genes (*CCL3* and *CCL3L1*) and cytokine genes (*IL1B*, *TNFSF10*, *MIF*, and *HGF*), which are involved in inflammatory signaling and NF-_K_B–mediated cell survival, were expressed by CMML monocytes, DCs, pDCs, and MyHPCs, whereas *IFNG* was only upregulated in pDCs. These populations also expressed the receptors of these ligands (i.e., *CCR1*, *CCR5*, *ADRB2*, *TNFRSF10B*, *CD74*, *CD44,* and type II *IFNR1*, respectively), which suggests that an aberrant feedback loop among different cell types contributes to CMML maintenance.

CMML monocytes, pDCs, and DCs also gained cell-to-cell interactions with NK and eCD8T cells. Interactions involving the HLA-E–KLRC1/2, BAG6–NCR3, CD48–CD244, and LGALS9–HAVCR2 ligand–receptor pairs were the most common (Supplementary Fig. S2H). This observation has added significance in light of recent findings that HLA-E expression by different cell types, including monocytes, inhibits NK cells’ and T cells’ release of IFNγ and compromises their functions through its interaction with the CD94/NKG2A inhibitory receptors^21-23^. Consistent with these data, both CMML NK cells (cluster 8) and eCD8T cells (cluster 3) had increased expression levels of immune checkpoint genes associated with these cells’ exhaustion (e.g., *KLRG1*, *TIGIT, LAG3,* and *CD160*) compared with those from HDs (Supplementary Fig. S2I)^24-26^.

Taken together, these data suggest that *RAS* pathway**–**mutated CMML HSPCs and downstream My/Mo cells undergo significant transcriptional rewiring that maintains these cells’ proliferation and suppresses the immune microenvironment, thus enabling immune escape and clonal expansion through cell-intrinsic and -extrinsic inflammatory networks.

### *RAS* pathway–mutated HSCs undergo epigenetic reprogramming and drive CMML BP after HMA therapy failure

To evaluate the cellular and molecular dynamics of CMML progression, we performed scRNA-seq analysis of Lin^−^CD34^+^ HSPCs isolated from BM samples longitudinally obtained from 5 CMML patients at diagnosis and BP (Fig. 3A; Supplementary Fig. S3A; Supplementary Table S7). HSPCs isolated from BM samples obtained at BP maintained aberrant differentiation towards the My/Mo lineage (Fig. 3B) and had upregulated genes belonging to the NF-_K_B signaling pathway (Fig. 3C). Importantly, *MCL1*, an anti-apoptotic member of the BCL2 family and a downstream effector of NF-_K_B pathway, was significantly upregulated in HSCs (cluster 6) and eMyHPCs (clusters 0 and 1) at BP compared with those at diagnosis (Supplementary Fig. S3B).

**Figure 3.**
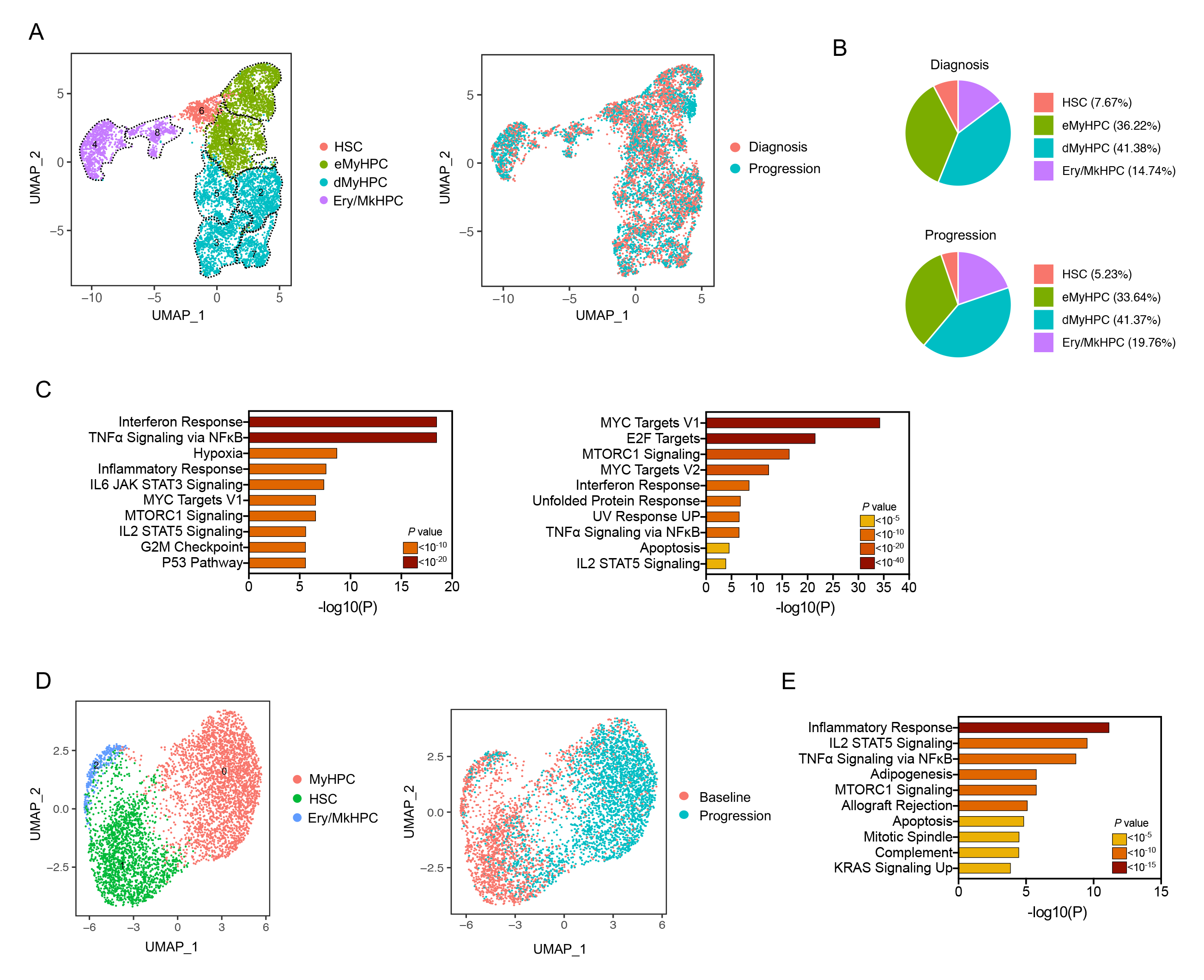
*RAS* pathway–mutated HSCs undergo epigenetic reprogramming and drive CMML BP after HMA therapy failure. (A) UMAP of scRNA-seq data for pooled single Lin^−^CD34^+^ cells isolated from BM samples of 5 CMML patients at diagnosis (n=1,840) and at BP after HMA therapy failure (n=1,711). Each dot represents one cell. Different colors represent the cluster cell type identity (left) or sample origin (right). HSC, hematopoietic stem cells; eMyHPC, early myeloid hematopoietic progenitor cells; dMyHPC, differentiated myeloid hematopoietic progenitor cells; Ery/MkHPC, erythroid/megakaryocyte hematopoietic progenitor cells. Dashed lines indicate single clusters in each cell type population. (B) Distribution of Lin^−^CD34^+^ cell types at diagnosis (top) and BP (bottom) among the clusters shown in Fig. 3A. (C) Pathway enrichment analysis of the genes that were significantly upregulated in HSCs (left) and dMyHPCs (right) at the time of BP after HMA therapy failure compared with those at diagnosis (adjusted *P* ≤ 0.05). The top 10 Hallmark gene sets are shown. (D) UMAP of scATAC-seq data for pooled Lin^−^CD34^+^ cells isolated from BM samples obtained from a CMML patient at diagnosis (n=2,027) and at BP after HMA therapy failure (n=2,895). Each dot represents one cell. Different colors represent the cluster identity (left) or sample of origin (right). HSC, hematopoietic stem cells; MyHPC, myeloid progenitor cells; Ery/MkHPC, erythroid/megakaryocyte hematopoietic progenitor cells. (E) Pathway enrichment analysis of genes whose distal elements were enriched in open chromatin regions in HSCs (cluster 1, shown in Fig. 3D) at the time of BP as compared with those at diagnosis (adjusted P ≤ 0.05). The top 10 Hallmark gene sets are shown.

To evaluate whether HSPCs’ transcriptional changes at BP were the result of epigenetic reprogramming in the more primitive HSCs, we performed single-cell assays for transposase-accessible chromatin with high-throughput sequencing (scATAC-seq) to profile the chromatin accessibility landscape in Lin^−^CD34^+^ HSPCs isolated from BM samples longitudinally obtained from a CMML patient at diagnosis or BP (whose BP was not associated with the clonal expansion of pre-existing or newly acquired *RAS* pathway– mutated clones, as shown in Supplementary Fig. S1A, S1B). Our analysis identified 3 clusters with distinct TF binding motif enrichment in the open chromatin regions (Fig. 3D; Supplementary Fig. S3C; Supplementary Table S8). MyHPCs (cluster 0) were characterized by open chromatin regions in the binding motifs of the myeloid TFs SPI1B and CEBPA and TFs belonging to the FOS and JUN families. HSCs (cluster 1) had the highest activities of TFs involved in stemness maintenance, such as those belonging to the nuclear retinoid receptor and EGR families of TFs. Ery/MkHPCs (cluster 2) were characterized by open chromatin regions in binding motifs for GATA TFs.

To determine whether CMML HSCs at BP were poised for myeloid differentiation, we analyzed the open chromatin peaks at the genes’ distal elements, which define cell identity and differentiation trajectories more precisely than promoter regions do^27^. This analysis showed that HSCs at BP had increased open chromatin peaks at the distal elements of genes involved in NF-_K_B pathway activation (Fig. 3E). Consistent with these results and our transcriptomic data, downstream MyHPCs had increased activity of TFs involved in NF- _K_B signaling activation, including FOS, FOSB, FOSL2, JUN, JUNB, NR4A2, FOSL1, and RORA (Supplementary Fig. S3D), and open chromatin peaks enriched at the promoters of these TFs’ target genes (Supplementary Fig. S3E), including *MCL1* (Supplementary Fig. S3F).

Together, these data suggest that *RAS* pathway–mutated CMML HSCs undergo epigenetic reprogramming at BP, and that their reprogramming drives MyHPCs’ transcriptional upregulation of NF-_K_B signaling–mediated anti-apoptotic pathways to maintain survival after HMA therapy failure.

### *RAS* pathway–mutated CMML cells rely on *MCL1* overexpression to maintain their survival at BP

To elucidate whether MyHPCs’ transcriptional and epigenetic reprogramming drives BP, we performed scRNA-seq analysis of MNCs isolated from sequential *RAS* pathway**–** mutated CMML BM samples obtained from 6 CMML patients at diagnosis and BP (Fig. 4A; Supplementary Fig. S4A; Supplementary Table S9). Compared with those at diagnosis, MNCs at BP had a significantly higher frequency of Lin^−^CD34^+^ MyHPCs (2.8% vs 22.4%, respectively; clusters 7, 8, and 14) (Fig. 4B; Supplementary Fig. S4B). These results were confirmed by flow cytometry analysis of Lin^−^CD34^+^ cells in 70% of the patients (Supplementary Fig. S4C), which suggests that HMA therapy failure is mostly driven by the expansion of the HSPC compartment. Consistent with our results in the Lin^−^CD34^+^ compartment, differential expression analysis confirmed that, compared with MyHPCs at diagnosis, MyHPCs at BP had upregulated genes involved in TNFα-mediated NF-_K_B activation (Supplementary Fig. S4D), including *MCL1* (Supplementary Fig. S4E). The upregulation of these genes was maintained in downstream My/Mo progenitors (My/MoPs; clusters 5 and 17) and was significantly increased in the monocytic populations (clusters 2, 3, 11, 15, and 20) (Fig. 4C; Supplementary Fig. S4F).

**Figure 4.**
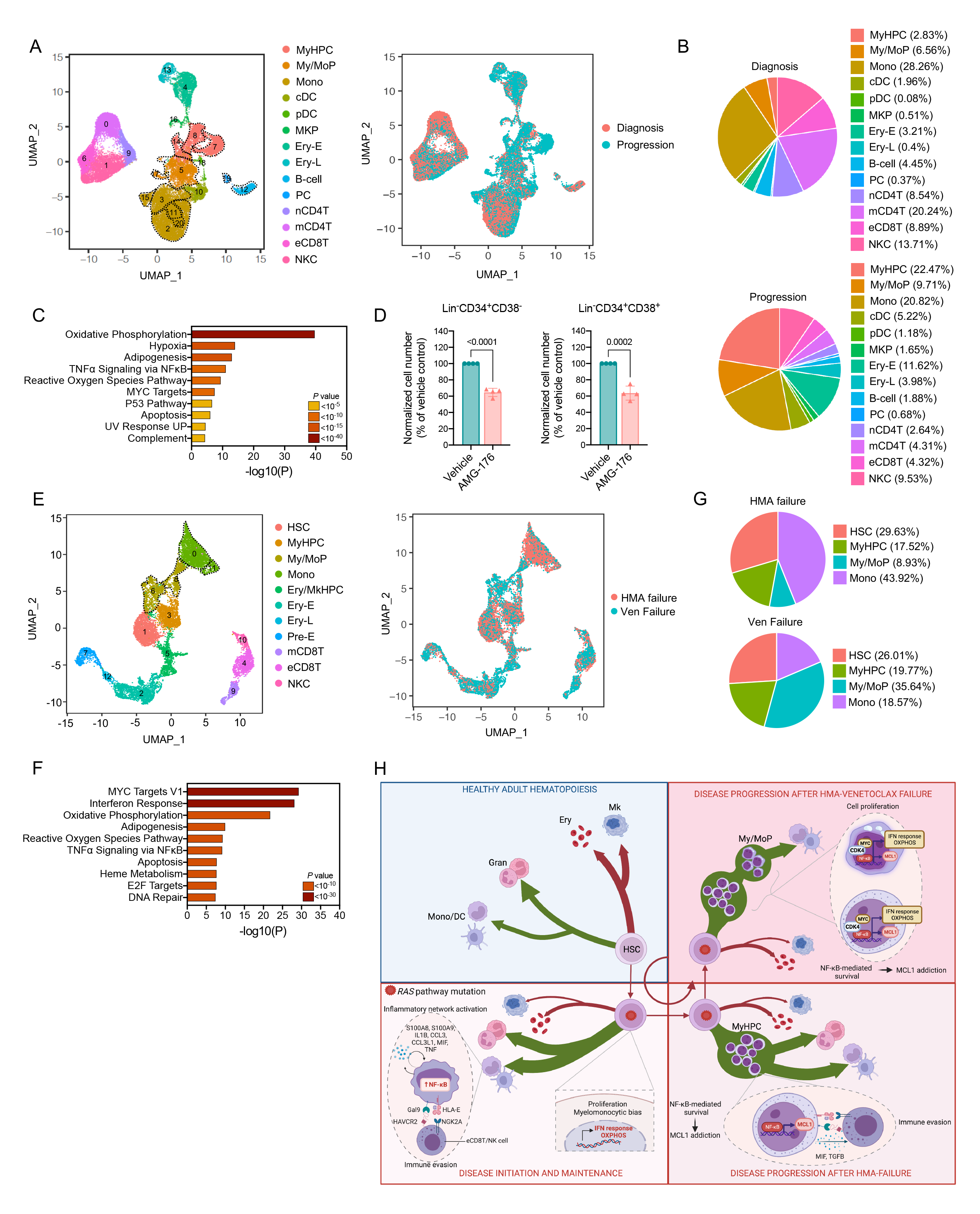
*RAS* pathway–mutated CMML cells rely on *MCL1* overexpression to maintain their survival at BP. (A) UMAP of scRNA-seq data for pooled single MNCs isolated from BM samples of 6 CMML patients at diagnosis (n=16,372) and at BP after HMA therapy failure (n=19,541). Each dot represents one cell. Different colors represent the cluster cell type identity (left) or sample of origin (right). MyHPC, myeloid hematopoietic progenitors; My/MoP, myelo/monocytic progenitors; Mono, monocytes; cDC, classical dendritic cells; pDC, plasmacytoid dendritic cells; MKP, megakaryocyte precursors; Ery-E, early erythroid precursors; Ery-L, late erythroid precursors; B-cell, B lymphocytes; PC, plasma cells; nCD4T, naïve CD4^+^ T cells; mCD4T, memory CD4^+^ T-cells; eCD8T, effector CD8 T-cells, NKC, natural killer cells. Dashed lines indicate single clusters in each cell type population. (B) Distribution of MNC types at diagnosis (top) and (bottom) among the clusters shown in Fig. 4A. (C) Pathway enrichment analysis of the genes that were significantly upregulated in the monocytic populations shown in Fig. 4A the time of BP after HMA therapy failure compared with those at the time of diagnosis (adjusted *P* ≤ 0.05). The top 10 Hallmark gene sets are shown. (D) Numbers of live Lin^−^CD34^+^CD38^−^ HSCs and Lin^−^CD34^+^CD38^+^ MyHPCs from CMML patients with BP after treatment with vehicle or 20 nM AMG-176 (n=4) for 48 h. Lines represent means ± SDs. Statistical significance was calculated using a two-tailed Student’s t- test (\*\*\**P*<0.001; \*\*\*\**P*<0.0001). (E) UMAP of scRNA-seq data for pooled single MNCs isolated from BM samples obtained from a representative CMML patient at the time of BP after HMA therapy failure (n=6,209) and after the failure of venetoclax-based therapy (n=6,795). Each dot represents one cell. Different colors represent the cluster cell type identity (left) or the sample of origin (right). HSC, hematopoietic stem cells; MyHPC, myeloid hematopoietic progenitor cells; My/MoP, myelo/monocytic progenitors; Mono, monocytes; Ery/MkHPC, erythroid/megakaryocytic hematopoietic progenitor cells; Ery-E, early erythroid precursors; Ery-L, late erythroid precursors; Pre-E, pre-erythrocytes; mCD8T, memory CD8^+^ T cells; eCD8T, effector CD8^+^ T cells; NKC, natural killer cells. (F) Pathway enrichment analysis of the genes that were significantly upregulated in MyHPCs at the time of venetoclax failure compared with those at the time of HMA therapy failure (adjusted *P* ≤ 0.05). The top 10 Hallmark gene sets are shown. (G) Distribution of myeloid cell types among the myeloid compartments at HMA therapy (top) and venetoclax-based therapy (bottom) failure. (H) Proposed working model of *RAS* pathway–mutated CMML initiation and progression after HMA and venetoclax-based therapies. Compared with physiological adult hematopoiesis (top left), *RAS* pathway–mutated CMML HSPCs undergo proliferation and monocytic differentiation in response to inflammatory responses while maintaining an intact apoptotic program. Inflammatory reprograming is exacerbated in downstream monocytic populations, which contributes to disease maintenance (bottom left). At BP after HMA therapy failure, *RAS* pathway–mutated CMML HSCs undergo epigenetic reprogramming and drive the expansion of downstream MyHPCs. MyHPCs and downstream monocytes rely on NF-_K_B signaling–mediated anti-apoptotic pathways to maintain survival and suppress the immune microenvironment (bottom right). NF-_K_B signaling–mediated survival pathway activation persists after venetoclax therapy and leads to treatment resistance and failure (top right).

Compared with MNCs at baseline, MNCs at BP exacerbated to a greater degree the cellular communication networks between cDCs, MyHPCs, My/MoPs, pDCs, monocytes, and eCD8T cells, mainly through immune-suppressive interactions between the HLA-E– KLRC1 and TGFB1–TGFBR3 ligand–receptor pairs (Supplementary Fig. S4G, S4H; Supplementary Table S10).

Together, these data suggest that *RAS* pathway–mutated CMML MyHPCs and monocytes rely on *MCL1*-driven anti-apoptotic pathways to maintain their survival and expand after therapy failure. To test this hypothesis, we treated Lin^−^CD34^+^ HSPCs isolated from the BM of patients with *RAS* pathway–mutated CMML with the MCL1 inhibitor AMG-176^28^ (at a dose that did not deplete Lin^−^CD34^+^CD38^−^ or Lin^−^CD34^+^CD38^+^ HSPCs isolated from the BM of HDs) in co-culture with mesenchymal stromal cells (Supplementary Fig. S4I). AMG-176 significantly decreased the numbers of Lin^−^CD34^+^CD38^−^ and Lin^−^ CD34^+^CD38^+^ HSPCs isolated from BM samples obtained from patients with *RAS* pathway– mutated CMML at BP (Fig. 4D). However, AMG-176 did not significantly affect the survival of HSPCs isolated from BM samples obtained from patients at diagnosis (Supplementary Fig. S4J), which confirms that CMML HSPCs maintain an intact apoptotic program at disease initiation.

Importantly, *BCL2* was not overexpressed in either CMML HSPCs or downstream My/Mo populations at BP after HMA therapy failure (Supplementary Fig. S4K), which is consistent with our previous clinical observation that CMML patients in whom HMA therapy has failed do not benefit from second-line therapy with venetoclax^10^. Indeed, scRNA-seq analysis of MNCs isolated BM samples longitudinally obtained from one representative CMML patient whose disease progressed after HMA therapy failure and did not respond to venetoclax therapy (Fig. 4E; Supplementary Fig. S4L; Supplementary Table S11) revealed that MyHPCs further exacerbate the expression of genes involved in TNFα-mediated NF-_K_B pathway activation (Fig. 4F), including *MCL1* (Supplementary Fig. S4M). Venetoclax failure was associated with a significant expansion of downstream My/MoPs in the myeloid compartment (Fig. 4G), and these cells also showed persistent high expression of *MCL1* (Supplementary Fig. S4M).

Together, our findings suggest that venetoclax therapy cannot overcome *RAS* pathway mutation–induced transcriptional reprogramming during My/Mo differentiation and provide a rationale for targeting the effectors of the NF-_K_B signaling pathway in patients with *RAS* pathway–mutated CMML.

## DISCUSSION

Whereas the dissection of the molecular landscape of CMML initiation and progression has significantly advanced our understanding of the pathogenesis of CMML,^1,29-32^ the development of more effective therapeutic approaches to improve patient survival has been delayed by our poor understanding of the ways in which genetic alterations affect distinct transcriptional states of My/Mo differentiation.

Mutations in *RAS* pathway genes, which are present in 30% of CMML patients^32^, are enriched during disease progression in up to 90% of cases, and predict a higher risk of and a shorter time to relapse after HMA and venetoclax therapy^33^. Currently, there are no other therapies that improve the survival duration of patients with *RAS* pathway–mutated CMML.

Using single-cell multi-omics technologies, we sought to dissect the biological mechanisms behind *RAS* pathway mutation–induced CMML evolution with the overall goal of identifying cellular vulnerabilities that could be therapeutically targeted to halt disease progression. We found that *RAS* pathway–mutated HSPCs at disease initiation significantly upregulate the NF-_K_B pathway–mediated inflammatory transcriptional signaling that drives these cells’ differentiation towards the My/Mo lineage while maintaining an intact apoptotic program. These HSPCs’ inflammatory reprogramming was exacerbated in downstream monocyte populations, which expressed high levels of cytokines and cell surface receptors involved in NF-_K_B pathway activation. These results suggest that disease initiation and maintenance rely on the activation of cell-intrinsic and -extrinsic inflammatory networks and provide a rationale for using inhibitors of NF-_K_B–associated inflammatory signaling cascades as a frontline treatment for patients with *RAS* pathway–mutated CMML. These findings have significant implications in light of the fact that several inflammation-targeting therapies currently in clinical development have shown great potential for treating patients with myeloid malignancies^34-36^.

Consistent with the long-standing observation that apoptosis inhibition contributes to therapy resistance and cancer progression, we found that *RAS* pathway–mutated CMML HSPCs isolated from BM samples obtained at BP depended on MCL1, an anti-apoptotic downstream effector of the NF-_K_B pathway, to maintain their survival and undergo clonal expansion. Targeting MCL1 activity with the small molecule AMG-176 significantly depleted these HSPCs, which supports the selective use of MCL1 inhibitors to treat patients with *RAS* pathway–mutated CMML in whom HMA therapy has failed. These results are in accordance with previous findings showing that *NRAS*-mutant monocytic subclones that emerge at AML relapse depend on MCL1, not BCL2, for energy production and survival ^37^ and explain why *RAS* pathway–mutated CMML is resistant to venetoclax, a selective BCL2 inhibitor^10^. Consistent with this observation, our scRNA-seq analysis of BM MNCs from one representative patient with venetoclax-resistant disease confirmed that BCL2 inhibition cannot overcome the activation of NF-_K_B pathway–mediated inflammatory and survival mechanisms in HSPCs and downstream My/Mo populations (Fig. 4H).

In conclusion, this study underscores the importance of dissecting the ways in which specific genetic drivers affect a cancer’s cell-of-origin to gain mechanistic insights into therapy failure and thereby develop selective therapeutic approaches to halt disease progression. Given that the *RAS* pathway mutation–induced reprogramming of CMML cells is a multi-step process that affects multiple biological signaling pathways (inflammation, apoptosis, immune escape, etc.) in distinct BM cell types, our findings also suggest that only combination therapies that target these pathways simultaneously are an effective approach to overcoming disease progression and prolonging the survival of patients with *RAS* pathway– mutated CMML.

## METHODS

### Human primary samples

BM aspirates were obtained from patients with *RAS* pathway–mutated CMML who were seen in MD Anderson’s Department of Leukemia. Samples were obtained with the approval of MD Anderson’s Institutional Review Board and in accordance with the Declaration of Helsinki. CMML diagnoses were assigned according to World Health Organization criteria^3^.

*RAS* pathway mutations were identified by targeted amplicon-based next-generation sequencing^38^. All available samples carrying *RAS* pathway mutations were included in the study. Baseline BM aspirates were collected from patients before any treatment. Sequential BM samples were collected after HMA or venetoclax therapy failure. The clinical characteristics of the patients with *RAS* pathway–mutated CMML are shown in Supplementary Tables S1 and S3. BM samples from HDs were obtained from AllCells (Alameda, CA) and MD Anderson’s Department of Stem Cell Transplantation. Written informed consent was obtained from all donors.

MNCs were collected from each BM sample immediately after BM aspiration using the standard gradient separation approach with Ficoll-Paque PLUS (catalog number #45-001- 752, Thermo Fisher Scientific). MNCs were cryopreserved and stored in liquid nitrogen until use. For cell sorting applications, MNCs were enriched in CD34^+^ cells using magnetic-activated cell sorting (MACS) with the CD34 Microbead Kit (catalog number #130-046-702, Miltenyi Biotec, Germany) and further purified by fluorescence-activated cell sorting (FACS) as described below.

### Clinical data analysis

Clinical datasets were analyzed using the SPSS 23.0 (SPSS, Inc., Chicago, IL, USA) and R (version 3.5.1) statistical software programs. Logistical regression analysis was performed using clinical, cytogenetic, and molecular characteristics in correlation with responses to HMA therapy. Associations between gene mutations and BP were assessed using data from 108 patients whose samples were sequenced using the 81-gene panel performed in the clinic. The dataset was randomly divided into a training set (30 patients with BP) and a testing set (5 patients with BP). A combination rule derived from selected features was trained using logistic regression in the training set and a fixed model in the testing set. Receiver operating characteristic (ROC) curves were generated using the “pROC” package in R (version 3.6.0). The 95% CIs for the areas under the ROC curves were estimated using the DeLong method^39^. The chi-square or Fisher exact test was used to analyze differences between categorical variables. Survival curves were generated using the Kaplan-Meier method and compared using log-rank tests. Responses to HMA- or venetoclax-based therapies were evaluated based on the International Working group 2003^40^ and 2006^41^ criteria for patients with secondary AML or CMML, respectively.

### FACS

Quantitative flow cytometry and FACS analyses of Lin^−^CD34^+^ cells were performed using previously described staining protocols^42,43^ and antibodies against: CD2, FITC, RPA- 2.10, BD Biosciences, 555326; CD3, FITC, SK7, BD Biosciences, 349201; CD4, FITC, S3.5, Thermo Fisher, MHCD0401; CD7, FITC, 6B7, BioLegend, 343104; CD11b, FITC, ICRF44, Thermo Fisher, 11-0118-42; CD14, FITC, MφP9, BD Biosciences, 347493; CD19, FITC, SJ25C1, BD Biosciences, 340409; CD20, FITC, 2H7, BD Biosciences, 555622; CD33, FITC, P67.6, Thermo Fisher, 11-0337-42; CD56, FITC, B159, BD Biosciences, 562794; CD235a, FITC, HIR2, BD Biosciences, 559943; CD34, BV421, 581, BD Biosciences, 562577; CD38, APC, HIT2, BioLegend, 303534, as we described previously^44^. Samples used for flow cytometry and FACS were acquired with a BD LSR Fortessa and a BD Influx Cell Sorter (BD Biosciences), respectively, and the cell populations were analyzed using FlowJo software (https://www.flowjo.com). All experiments included single-stained controls and were performed at MD Anderson’s South Campus Flow Cytometry & Cellular Imaging Facility.

### scRNA-seq analysis

ScRNA-seq analysis was performed as we described previously^44^. Briefly, live Lin^−^ CD34^+^ and MNC samples were enriched by FACS. Sample preparation and sequencing were performed at MD Anderson’s Advanced Technology Genomics Core. Sample concentration and cell suspension viability were evaluated using a Countess II FL Automated Cell Counter (Thermo Fisher Scientific). Samples were normalized for input onto the Chromium Single Cell A Chip Kit (10X Genomics), and single cells were lysed and barcoded for reverse transcription. Equal amounts of each uniquely indexed sample library were pooled together. Pooled libraries were sequenced using a NovaSeq6000 SP 100-cycle flow cell (Illumina). The Seurat package in R was used to analyze the digital expression matrix. Cells with fewer than 100 genes and fewer than 500 unique molecular identifiers detected were removed from further analysis. Principal component analysis and uniform manifold approximation and projection (UMAP) were used to reduce the dimensions of the data, and the first 2 dimensions were used in plots. To cluster the cells and determine the marker genes for each cluster, we used the FindClusters and FindAllMarkers functions, respectively. Differential expression analysis of the samples was performed using the FindMarkers function and the Wilcoxon rank-sum test. The Benjamini-Hochberg procedure was applied to adjust the false discovery rate. Functional enrichment analysis was performed using the Metascape software (https://metascape.org/gp/index.html#/main/step1)^45^. The human Hallmark set was used. Analyses were performed using gene annotation available in 2020-2022.

CellphoneDB (v2.0.0)^20^ was used to analyze the ligand–receptor interactions. Briefly, each cell type was separated by their disease classifications, and a separate run was performed for each disease classification. The connectome web was plotted using the igraph package in R.

The predicted upstream regulators of activated pathways were determined by Ingenuity Pathway Analysis (https://digitalinsights.qiagen.com/products-overview/discovery-insights-portfolio/analysis-and-visualization/qiagen-ipa). The top 15 TF, cytokine, and growth factor upstream regulators were plotted.

### scATAC-seq analysis

ScATAC-seq analysis was performed as we described previously^44^. Briefly, the scATAC Low Cell Input Nuclei Isolation protocol (10X Genomics) was used to isolate nuclei from FACS-purified cells. Extracted nuclei were used for the consecutive steps of the scATAC-seq library preparation protocol following 10X Genomics guidelines. Equal molar concentrations of uniquely indexed samples were pooled together. Pooled libraries were sequenced using a NextSeq500 150-cycle flow cell (Illumina). To identify specific TF activity for each cell cluster, we used the R package Seurat to analyze the TF-barcode matrix. Principal component analysis and UMAP were applied to reduce the dimensions of the data, and the first 2 dimensions were plottedCluster identity was determined based on the activity of master regulators of lineage commitment, as we^46^ and others^27,47^ described previously.

### scDNA and protein-seq analysis

Simultaneous analyses of DNA mutations and the cell-surface immunophenotype (scDNA and protein-seq) were performed as we described previously^48^ and according to the Mission Bio protocol using the custom-designed 37-gene myeloid panel kit and 48 oligo-conjugated antibodies against all major BM cell types (Biolegend). Briefly, cryopreserved BM MNCs were thawed, quantified, and then stained with the pool of the oligo-conjugated antibodies. Stained cells were washed and loaded onto the Tapestri^®^ machine for single-cell encapsulation, lysis, and barcoding. DNA libraries were extracted from the droplets followed by the purification using Ampure XP beads (Beckman Coulter). The supernatant from Ampure XP beads incubation contained antibody-tagged libraries was incubated with biotinated oligo (Integrated DNA Technologies) to capture the antibody tags, followed by the purification using streptavidin beads (Thermo Fisher Scientific). Purified DNA and antibody-tagged libraries were indexed and then sequenced on the Illumina NovaSeq 6000 or NextSeq 500 systems with 150 bp paired-end multiplexed runs.

The resulting files containing DNA and protein data were visualized using the Mission Bio Mosaic library version 1.8. Only manually curated and whitelisted variants were used. Variants were filtered using the below setting: min_dp=5, min_gq=0, min_vaf=21, max_vaf=100, min_prct_cells=0, min_mut_prct_cells=0, and min_std=0. Protein reads were normalized by centered log ratio, and subsequently underwent dimensionality reduction and clustering using Mosaic ‘run_pca’ (components=15), ‘run_umap’ (attribute=’pca’, n_neighbors=20, metric=’cosine’, min_dist=0), and ‘cluster’ (attribute=’umap’, method=’graph-community’, k= 150). Default parameters were used unless otherwise specified, and randomness was controlled in all steps. Heatmaps were separately visualized in R using the ComplexHeatmap package (https://bioconductor.org/packages/release/bioc/html/ComplexHeatmap.html).

### Primary cell culture assays

FACS-purified Lin^−^CD34^+^ HSPCs were resuspended in cytokine-free sterile RPMI medium supplemented with 10% FBS, 1% penicillin–streptomycin, and 0.1% amphotericin B and plated in 48-well plates previously seeded with low-passage (P ≤4) healthy BM-derived human mesenchymal cells. Co-cultures were incubated at 37 °C in a 5% CO_2_ atmosphere. After treatment with vehicle or AMG-176 (20 nM) for 48 h, cells were harvested and stained for quantitative flow cytometric analysis using the antibody panel described above and with AccuCheck Counting Beads (Thermo Fisher Scientific) added to each tube.

### Statistical analysis

Flow cytometry data were analyzed with Prism 8 software (https://www.graphpad.com). The figure legends indicate the statistical test(s) used in each experiment. Statistical significance was represented as \**P*<0.05, \*\**P*<0.01, \*\*\**P*<0.001, and \*\*\*\**P*<0.0001. All analyses involving human samples, investigators were blinded to sample annotations and patient outcomes. For replicated experiments, the number of replicates is indicated in the figure legends. No statistical method was used to predetermine sample size. No data were excluded from the analyses. The experiments were not randomized.

The graphical abstract was made using Biorender.com.

### Data availability

Datasets generated by scRNA-seq and ATAC-seq are accessible at GEO under accession number GSE218390.

## AUTHORS’ CONTRIBUTIONS

S.C., I.G.-G., H.Y., N.T., V.A., B.W., and P.L. performed the experiments; F.M. analyzed scRNA and scATAC-seq data; R.K.-S. analyzed targeted DNA sequencing data; Y.J.K, T.J.K., and K.T. performed the scDNA and protein-seq experiments and analyzed their data; K.C.-D. analyzed the flow cytometry data. G.M.-B identified the clinical samples included in the study. G.M.-B. and F.D. analyzed the clinical data; G.A.-A. provided the HD samples; J.J. R.-S., H.K. and G.G.-M. made critical intellectual contributions throughout the study. S.C. designed the research and wrote the manuscript.

## ACKNOWLEDGMENTS

This work was supported by philanthropic contributions to MD Anderson’s AML/MDS Moon Shot, by the Edward P. Evans Foundation, by the National Institutes of Health through MD Anderson’s Leukemia SPORE grant (P50 CA100632), and by a grant from the Cancer Prevention and Research Institute of Texas (RP140500). S.C. is a Scholar of the Leukemia and Lymphoma Society. This work used MD Anderson’s South Campus Flow Cytometry & Cellular Imaging Facility and the Advanced Technology Genomics Core, which are supported in part by the National Institutes of Health (National Cancer Institute) through MD Anderson’s Cancer Center Support Grant (P30 CA16672). The authors thank Joseph Munch for assistance with manuscript editing.

## SUPPLEMENTARY FIGURE LEGENDS

**Figure S1.**
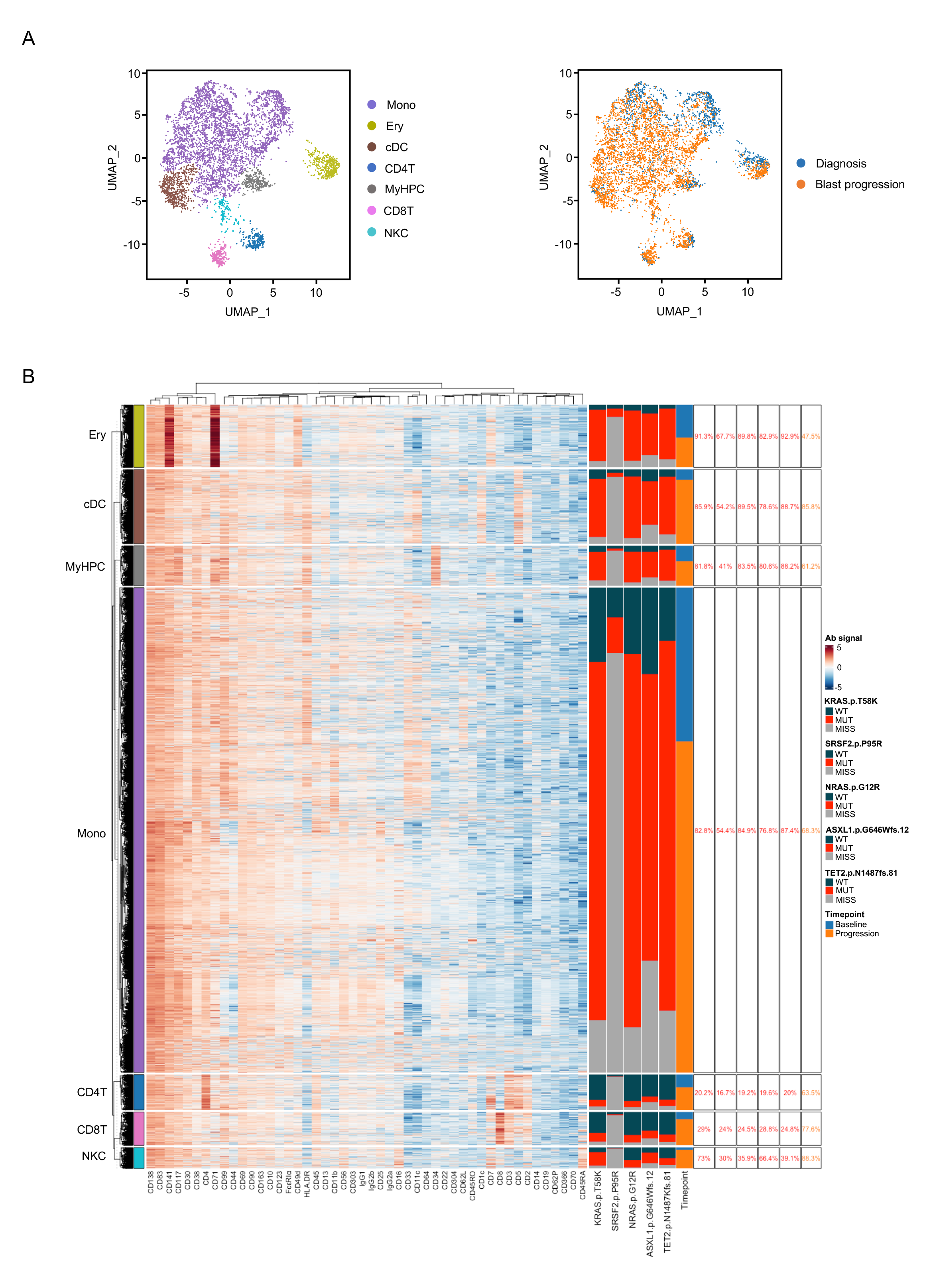
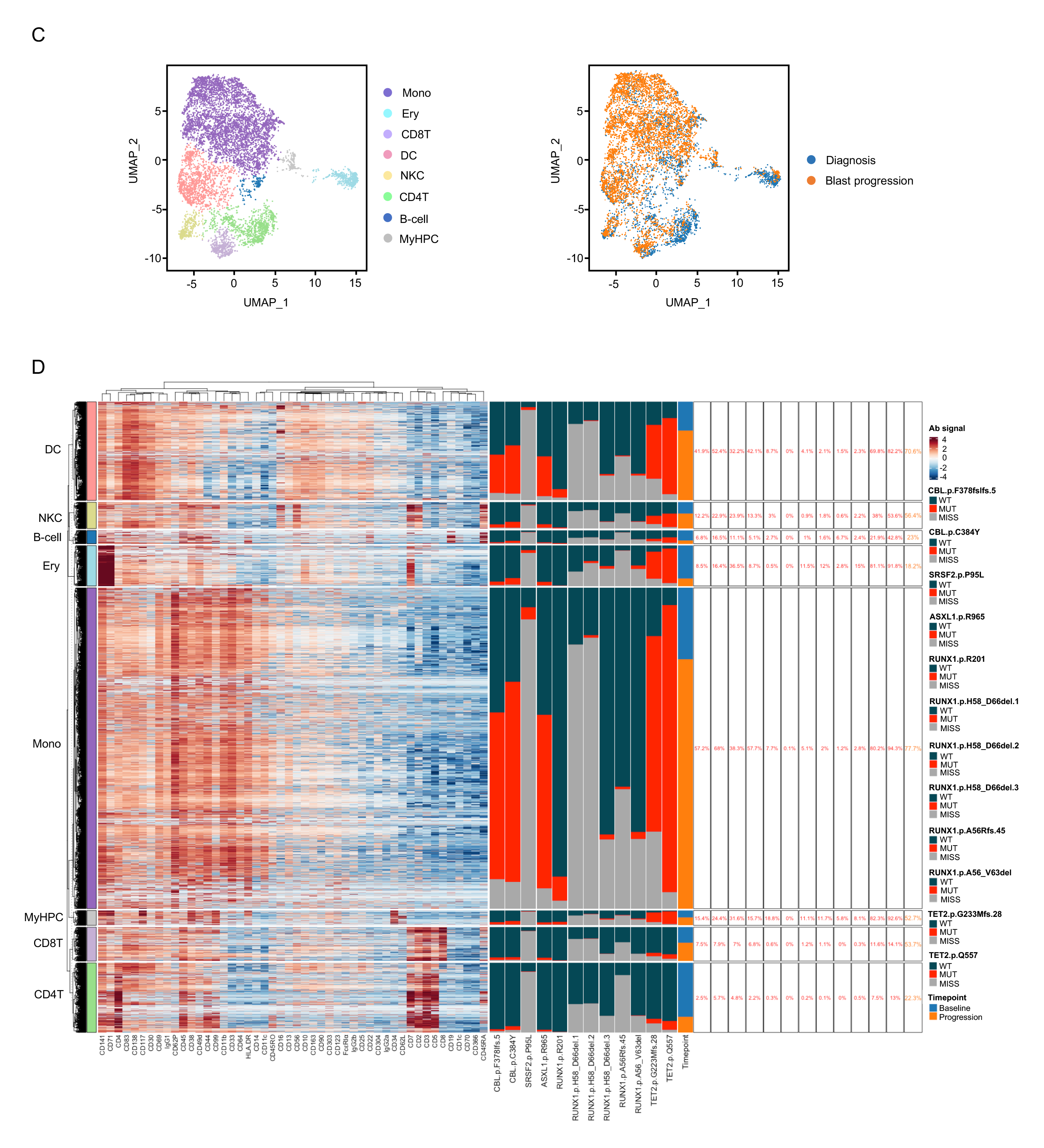
Mutations in *RAS* pathway signaling genes predict a high risk of CMML BP after HMA therapy failure. A) UMAP of scDNA and protein-seq data for pooled MNCs isolated from BM samples obtained from a CMML patient with pre-existing *KRAS^T58K^* and *NRAS^G12R^*mutations at diagnosis (n=1,826) and at BP after HMA therapy failure (n=4,001). BP was not associated with the clonal evolution of these mutations as they both had a VAF of approximately 50% at the onset of the disease. Each dot represents one cell. Cells are clustered based on immunophenotypic markers. Different colors represent cluster identity (left) or origin (right). Mono, monocytes; Ery, erythroblasts; cDC, classical dendritic cells; CD4T, CD4^+^ T-cells; MyHPC, myeloid hematopoietic progenitor cells; CD8T, CD8^+^ T-cells; NKC, natural killer cells. (B) Heatmap displaying DNA and protein reads from each sequenced cell type shown in Fig. S1A. Colors for protein data correspond to antibody-oligonucleotide intensity signals. High protein expression is depicted in red and low protein expression is depicted in blue. DNA colors correspond to the genotypes for each individual mutation per cell read (wild-type=dark grey, mutant=red, missing=light grey) based on cluster. Percentages correspond to the frequencies of mutant reads within each cluster for a given mutation. C) UMAP of scDNA and protein-seq data for pooled MNCs isolated from BM samples obtained from a CMML patient at diagnosis (n=3,213) and at BP after HMA therapy failure (n=5,342). BP was associated with the clonal evolution of a pre-existing *CBL^F378Ifs^* mutation and the acquisition of a previously undetected *CBL^C384Y^* mutation. Each dot represents one cell. Cells are clustered based on immunophenotypic markers. Different colors represent cluster identity (left) or origin (right). Mono, monocytes; Ery, erythroblasts; DC, classical dendritic cells; CD4T, CD4^+^ T-cells; B-cell, B lymphocytes, myeloid hematopoietic progenitor cells; CD8T, CD8^+^ T-cells; NKC, natural killer cells. (D) Heatmap displaying DNA and protein reads from each sequenced cell type as shown in Fig. S1C. Colors for protein data correspond to antibody-oligonucleotide intensity signals. Red indicates high protein expression, and blue indicates low protein expression. Colors for DNA data correspond to the genotype for each individual mutation per cell read (dark grey, wild type; red, mutant; light grey, missing) based on cluster. Percentages correspond to the frequencies of mutant reads within each cluster for a given mutation.

**Figure S2.**
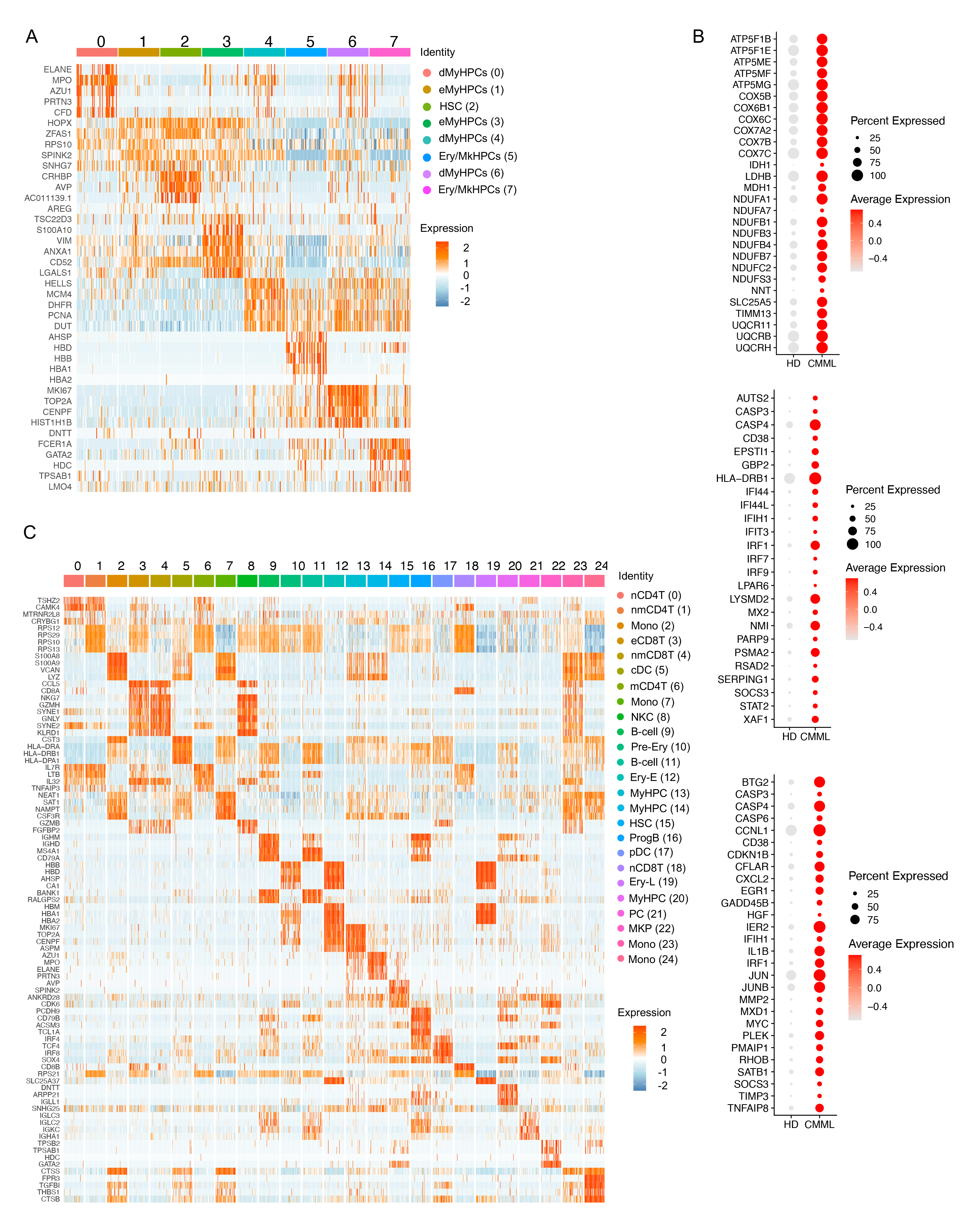
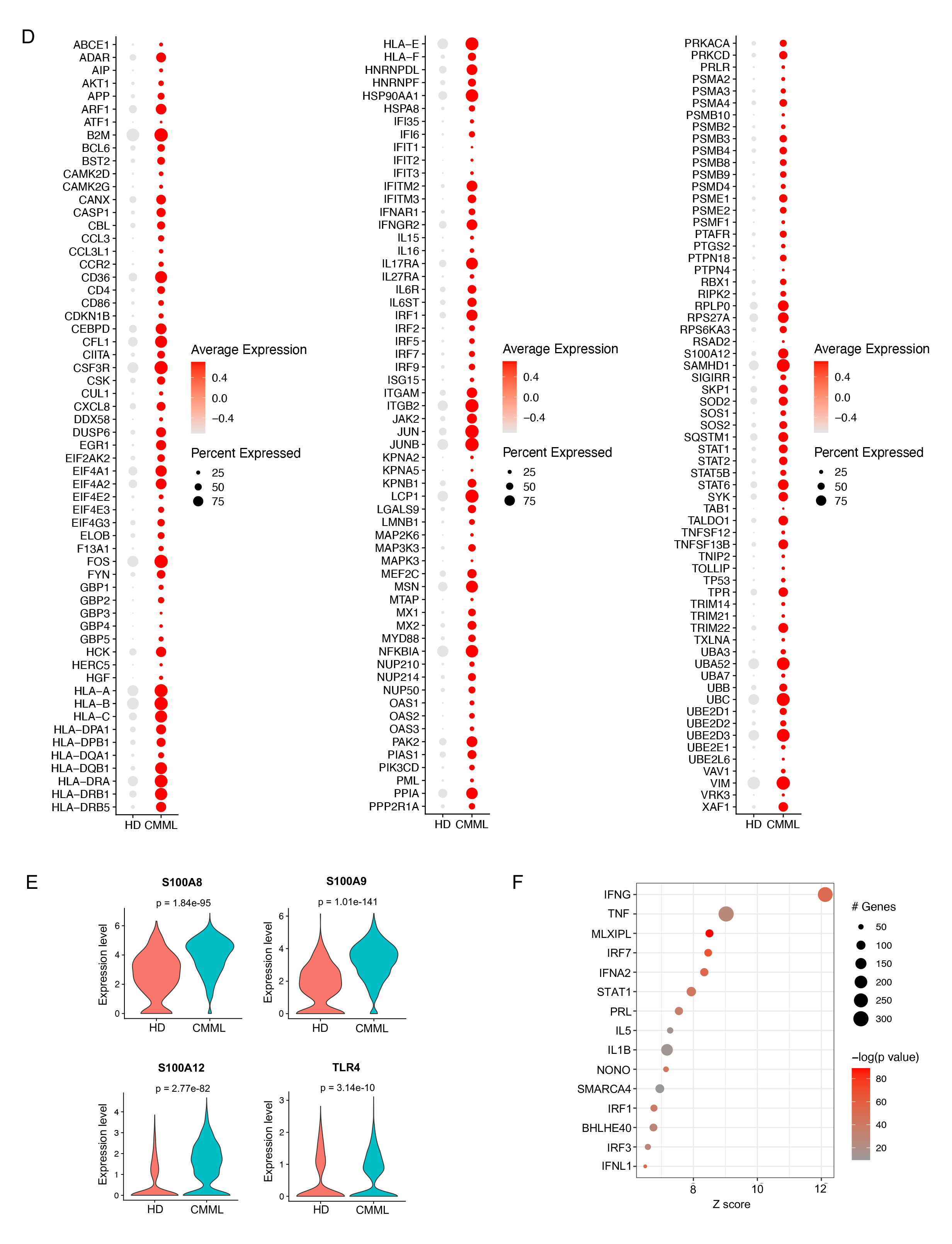
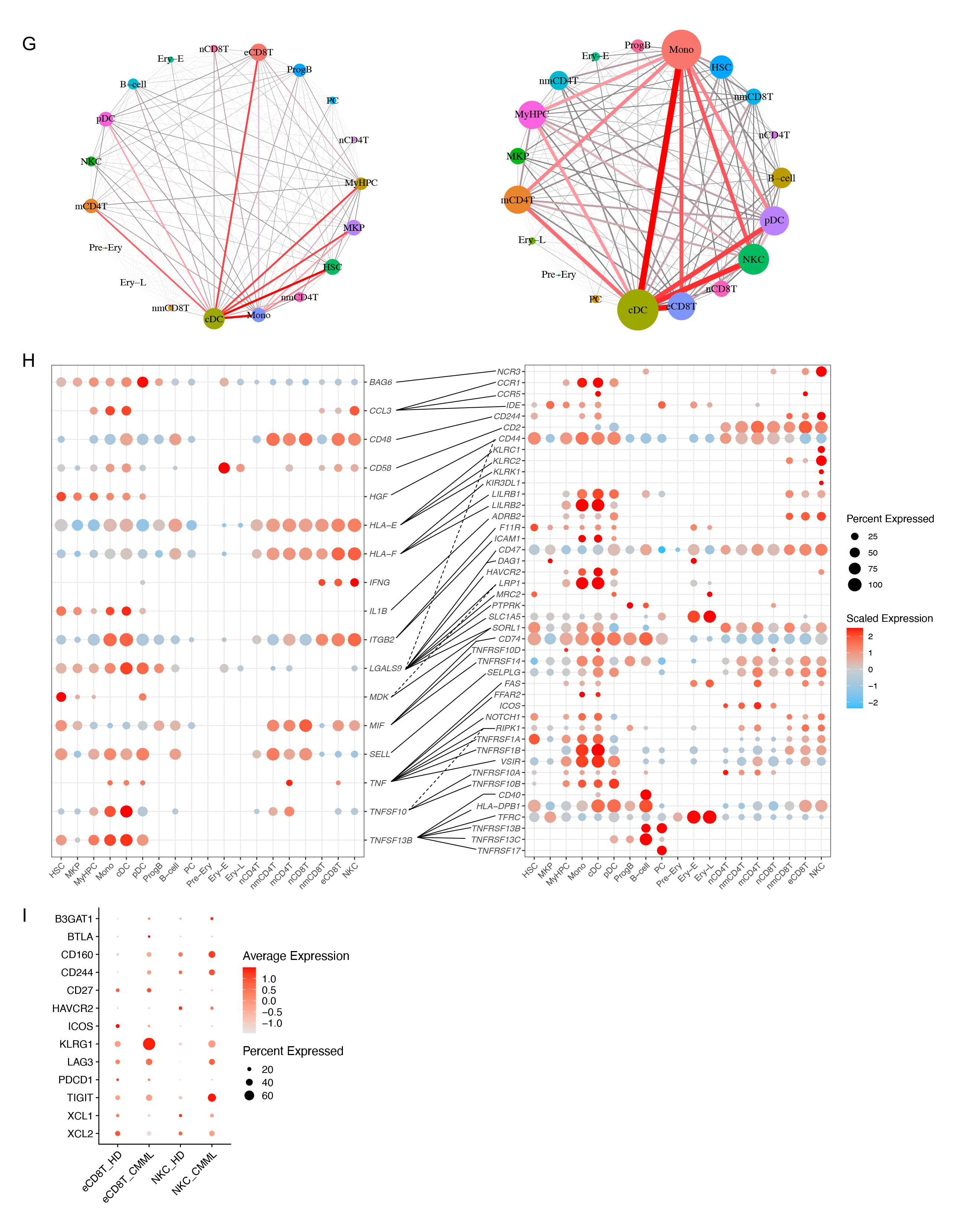
*RAS* pathway–mutated CMML cells activate cell-intrinsic and -extrinsic inflammatory networks. (A) Heatmap of the expression levels of the top 5 genes enriched in each of the 8 clusters shown in Fig. 2A. (B) Dot plots of the genes belonging to oxidative phosphorylation (top), IFN response (middle), and apoptosis (bottom) pathways that were significantly overexpressed in the CMML HSCs shown in Fig. 2A compared with those in HD HSCs. (C) Heatmap of the expression levels of the top 5 genes enriched in each of the 25 clusters shown in Fig. 2D. (D) Dot plots of the genes belonging to IFN response (left and middle) and NF-κB signaling (right) pathways that were significantly overexpressed in the CMML monocytes shown in Fig. 2D compared with those in HD monocytes. (E) Violin plots of expression levels of *S100A8, S100A9, S100A12,* and *TLR4* in CMML monocytes compared with those in HD monocytes (adjusted *P* = 1.84 × 10^−95^, 1.01 × 10^−141^, 2.77 × 10^−82^, and 3.14 × 10^−10^, respectively). (F) Dot plot showing the top 15 predicted upstream regulators of upregulated genes in CMML monocytes compared with HD monocytes. The color scale represents the –log_10_ *P*- value from the enrichment analysis. The size of the dot represents the number of downstream genes of each upstream regulator. The Z score represents the activation level of the upstream regulator. (G) Connectome web analysis of interacting cell types among MNCs isolated from HD (left) and CMML (right) BM samples. The vertex (i.e., colored cell node) size is proportional to the number of interactions to and from each cell type, and the thickness of each connecting line is proportional to the number of interactions between 2 nodes. (H) Dot plots showing the most significant ligand-to-receptor interactions that were gained in MNCs from CMML patients compared with those from HDs. (I) Dot plot of exhaustion markers in HD and CMML effector CD8^+^ T cells (eCD8T) and natural killer cells (NKC).

**Figure S3.**
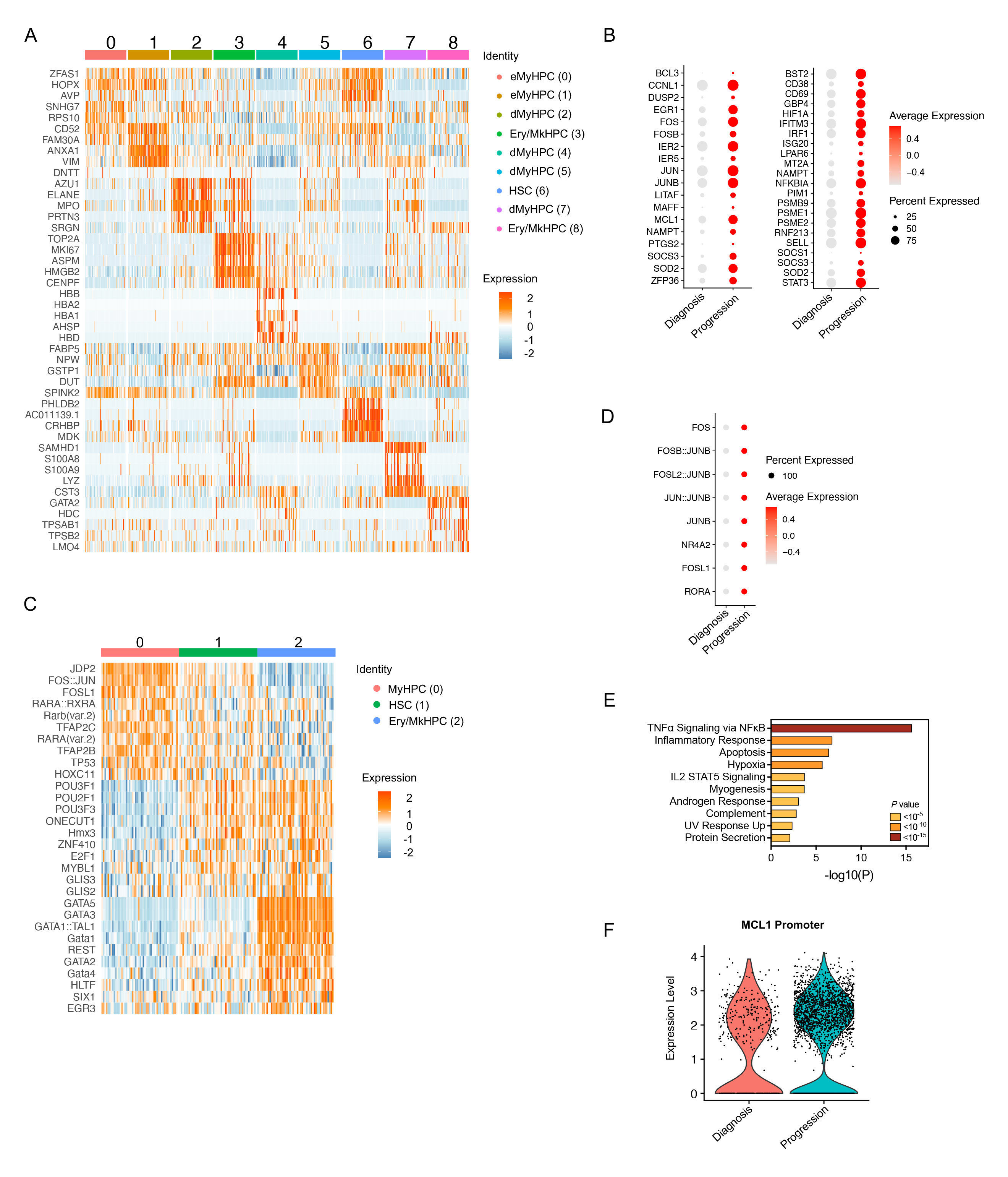
*RAS* pathway–mutated HSCs undergo epigenetic reprogramming and drive CMML BP after HMA therapy failure. (A) Heatmap of the expression levels of the top 5 genes enriched in each of the 9 clusters shown in Fig. 3A. (B) Dot plots of genes belonging to the NF-κB signaling pathway that were significantly upregulated in CMML HSCs (left) and eMyHPCs (right) at BP compared with those at diagnosis (adjusted P ≤ 0.05). (C) Heatmap of the activity of the top 5 TFs enriched in each of the 3 clusters shown in Fig. 3D. (D) Dot plot of the activity levels of the TFs involved in the NF-_K_B signaling pathway whose binding sites were significantly enriched in the open chromatin regions of CMML MyHPCs at BP compared with those at diagnosis (adjusted P ≤ 0.05). (E) Pathway enrichment analysis of genes whose promotors were enriched in accessible binding sites in CMML HSCs at BP compared with those in CMML HSCs at diagnosis (adjusted *P* ≤ 0.05). The top 10 Hallmark gene sets are shown. (F) Violin plots of open chromatin peak levels at the promoter of *MCL1* in CMML HSCs at BP compared with those in CMML HSCs at diagnosis (adjusted *P* = 3.03 × 10^−5^).

**Figure S4.**
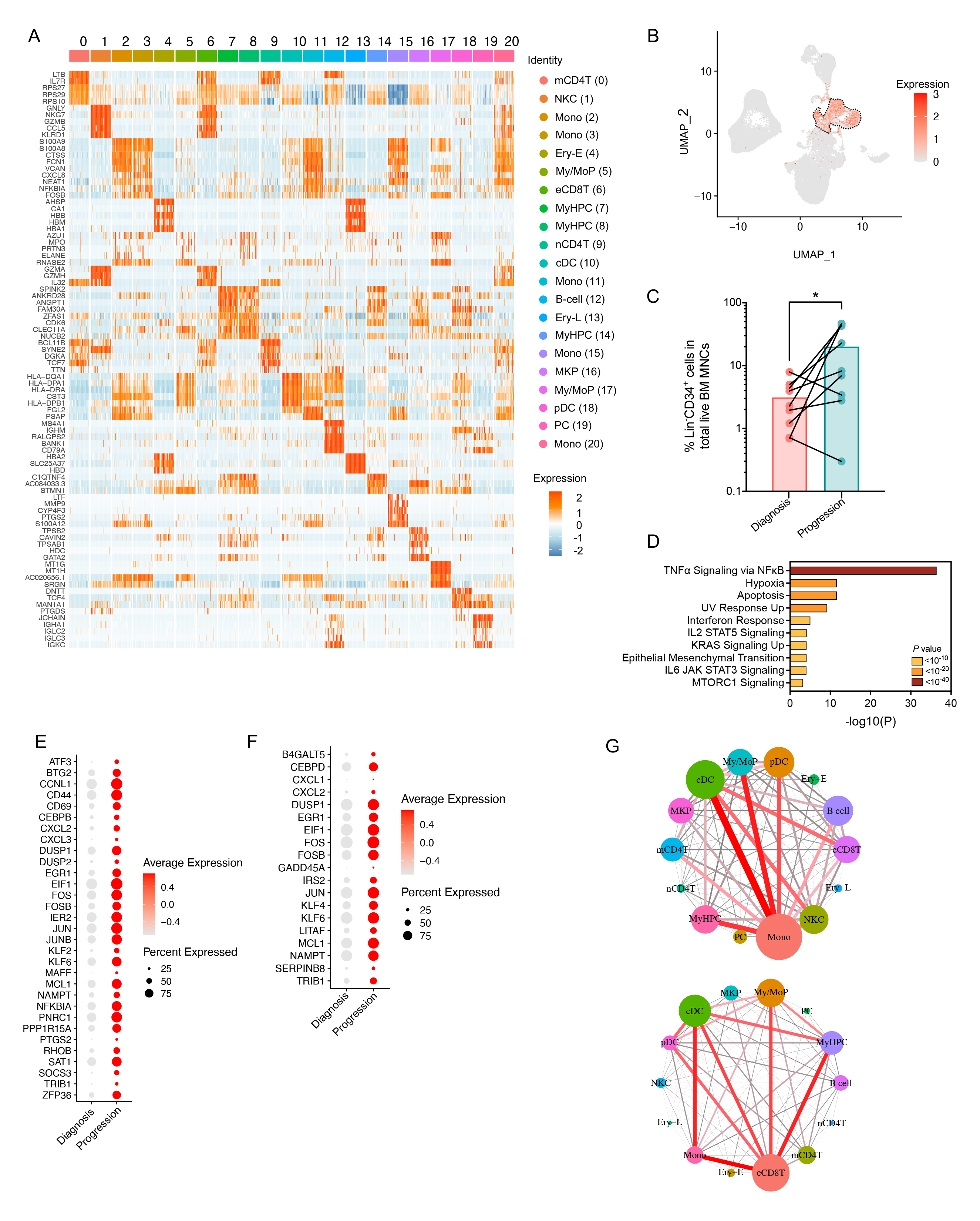
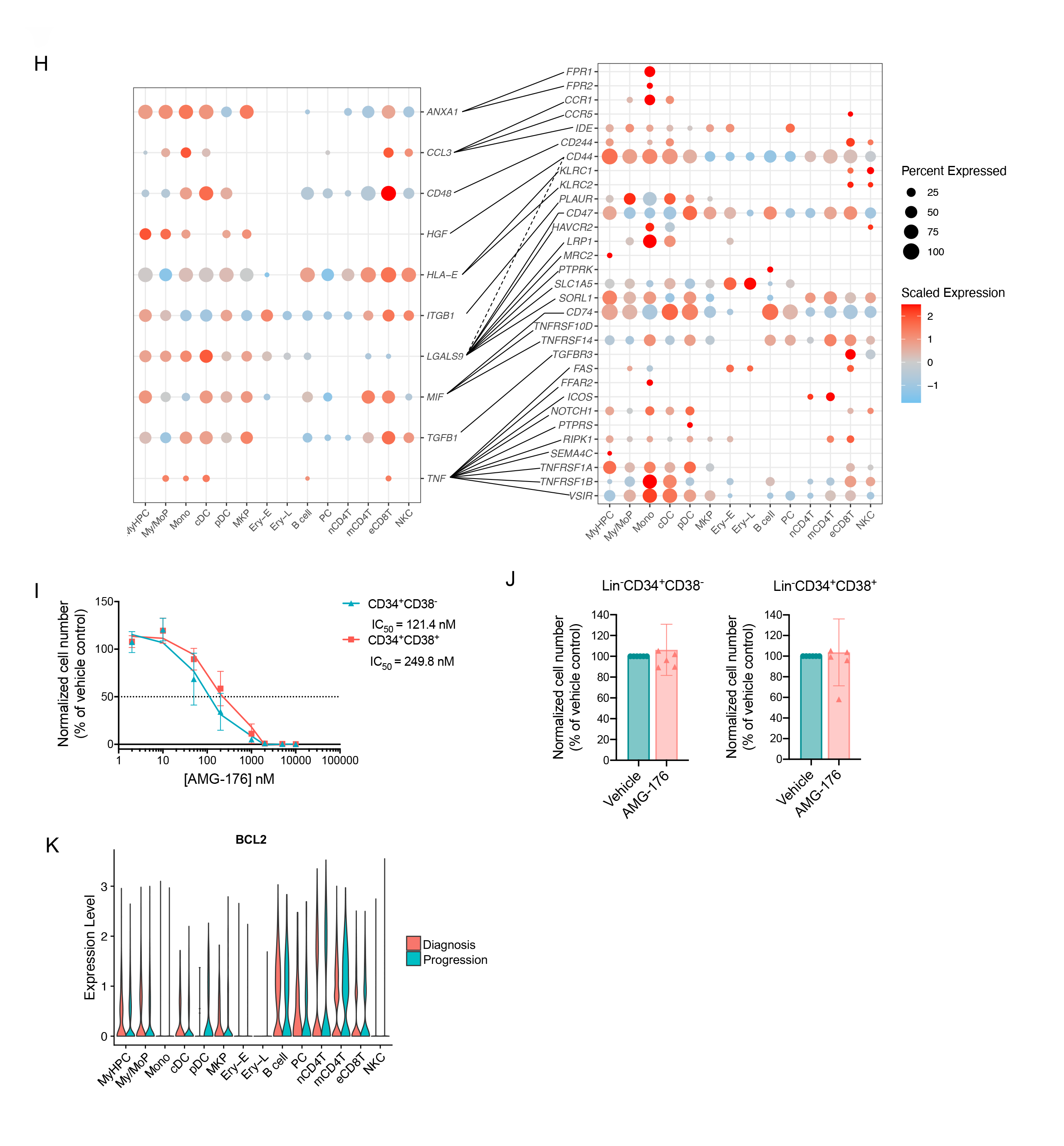
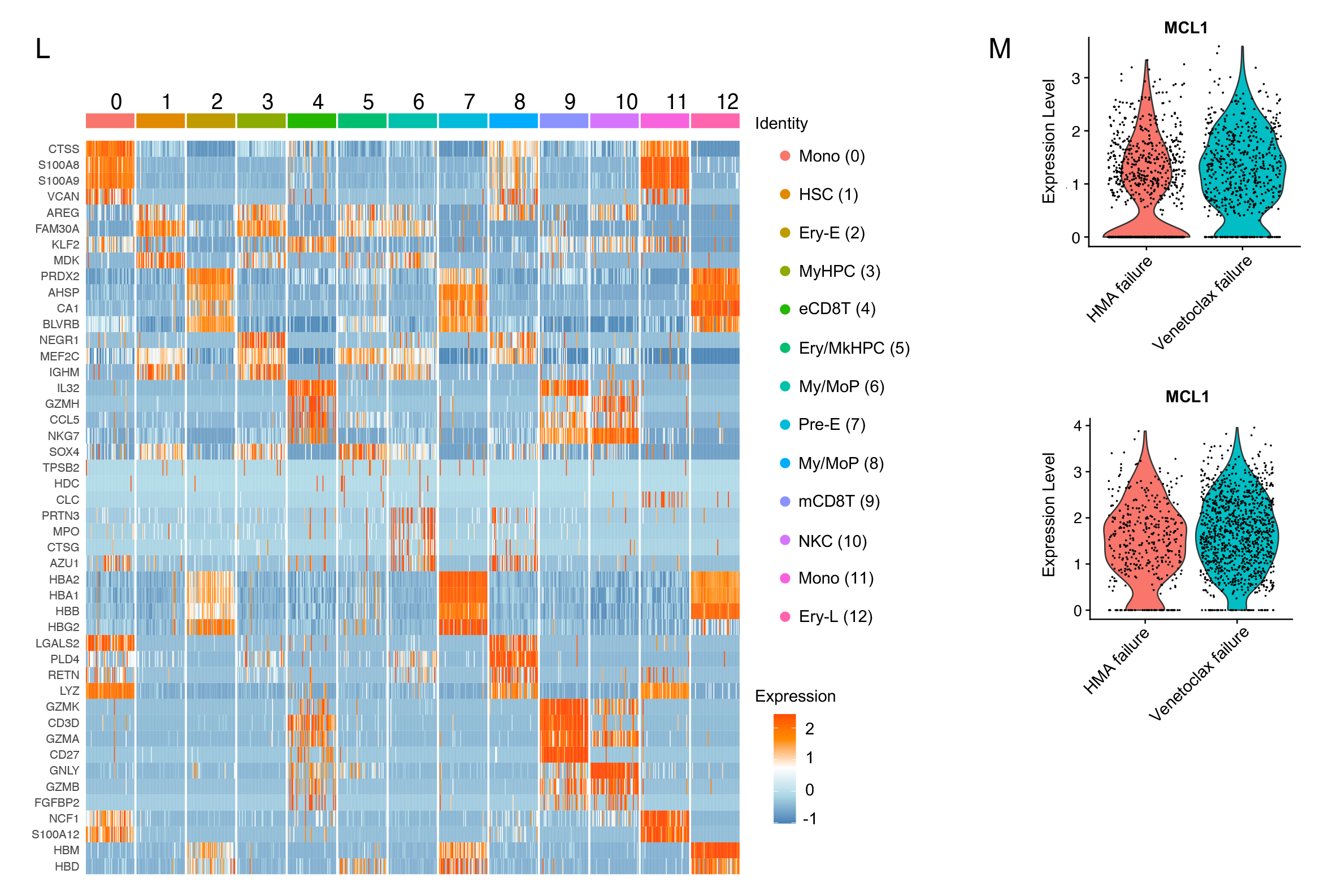
*RAS* pathway–mutated CMML cells rely on *MCL1* overexpression to maintain their survival at BP. (A) Heatmap of the expression levels of the top 5 genes enriched in each of the 21 clusters shown in Fig. 4A. (B) UMAP showing the distribution of *CD34* expression levels across the clusters shown in Fig. 4A. Red shading indicates normalized gene expression. Dashed lines indicate MyHPCs. (C) Frequency of Lin^−^CD34^+^ cells in MNCs from CMML BM samples sequentially collected at diagnosis and at BP after HMA therapy failure (n=9). Statistical significance was calculated using a paired two-tailed Student’s t-test (\**P* <0.05). (D) Pathway enrichment analysis of the genes that were significantly upregulated in MyHPCs at the time of BP compared with those at diagnosis (adjusted *P* ≤ 0.05). The top 10 Hallmark gene sets are shown. (E) Dot plots of genes belonging to the NF-κB signaling pathway that were significantly upregulated in CMML MyHPCs at BP compared with those at diagnosis (adjusted *P* ≤ 0.05). (F) Dot plots of genes belonging to the NF-κB signaling pathway that were significantly upregulated in CMML monocytes at BP compared with those at diagnosis (adjusted *P* ≤ 0.05). (G) Connectome web analysis of interacting cell types among BM MNCs from CMML patients at diagnosis (up) and BP (down). The vertex (i.e., colored cell node) size is proportional to the number of interactions to and from each cell type, and the thickness of each connecting line is proportional to the number of interactions between 2 nodes. (H) Dot plots showing the most significant ligand-to-receptor interactions that were gained in MNCs from CMML patients at diagnosis compared with those at BP (adjusted *P* ≤ 0.05). (I) Numbers of live cultured Lin^−^CD34^+^CD38^−^ and Lin^−^CD34^+^CD38^+^ cells from HD BM samples (n=2) after 48 h of treatment with AMG-176. Lines represent means ± SEMs. (J) Numbers of live Lin^−^CD34^+^CD38^−^ HSCs and Lin^−^CD34^+^CD38^+^ MyHPCs from CMML patients at diagnosis and after treatment with vehicle or 20 nM AMG-176 (n=6) for 48 h. Lines represent means ± SDs. A paired two-tailed Student’s t-test revealed no significant differences. (K) Violin plots of *BCL2* expression levels across each CMML MNC type at diagnosis compared with those at BP (no significant difference was detected). (L) Heatmap of the expression levels of the top 5 genes enriched in each of the 13 clusters shown in Fig. 4E. (M) Violin plots of *MCL1* expression levels in MyHPCs (top) and My/MoPs (bottom) at the time of HMA therapy failure compared with those at venetoclax failure (adjusted *P* = 1.12 × 10^−15^ and no significant difference, respectively)

